# The Martini 3 Metabolome

**DOI:** 10.64898/2026.03.06.710121

**Authors:** Christopher Brasnett, Chelsea M. Brown, Linus Grünewald, Jan A. Stevens, Siewert J. Marrink

## Abstract

Metabolites are ubiquitous in all living cells and are essential mediators of biochemical processes, serving either as substrates or as cofactors to enable the reactions. Capturing this diversity in computational workflows is important for allowing realistic simulations of the cytoplasm. Coarse-grained molecular dynamics enables the simulation of large scale systems up to the level of whole-cells, but is limited by the availability of refined parameters for all possible components in the system. In this work, we describe the parameterization of 186 common metabolites found in bacteria and eukaryotes within the framework of the Martini 3 force field. To showcase the behavior of Martini metabolites in a biological setting, we report simulations of protein-ligand binding and membrane permeation. The establishment of a Martini metabolome enables high-throughput simulations of metabolites interacting with other biomolecules, and opens the way for simulations of realistic cellular environments.

## Introduction

Molecular dynamics (MD) simulations have emerged as a powerful computational approach to investigate biological systems with spatio-temporal resolution nearly impossible to achieve in traditional laboratory settings. These simulations can provide detailed insights into how the full complexity of cellular environments influences molecular behavior, serving as a computational microscope for biological processes. Coarse-grained (CG) models offer access to larger systems and timescales by simplifying the representation. One of the most popular CG force fields is Martini, which broadly uses a 4-to-1 heavy atom to particle mapping, sufficient to retain chemical specificity^1^. While the latest version of the force field (Martini 3) has good coverage for simulating biomolecular systems, from lipids, proteins, and sugars to small organic molecules, extensive parameters for metabolites such as cofactors and nucleotides are still missing. Some metabolites have been modelled in previous iterations of the force field^2–4^, and there is a small selection compatible with the current version^1,5–8^, but the vast number of metabolites found *in vivo* are not yet captured in the Martini framework.

Simulations are increasingly moving towards biological levels of accuracy in terms of their components^9–12^, attempting to reach an *in situ* composition^13^. One of the major bottlenecks in assembling systems such as these is the availability of parameters for all components in the system. This challenge is particularly evident for metabolites, where the vast chemical diversity of molecules requires extensive parameterization efforts. The magnitude of this problem becomes especially clear when modeling whole-cells like the JCVI-Syn3A minimal cell^10^. Despite the reduced complexity of this synthetic organism, its metabolome contains around 200 distinct molecules^14,15^. Equally this is a problem for modelling organelles such as the mitochondria which have over 100 metabolites reported in the matrix alone^11,16,17^. The majority of the metabolites from both systems are not yet available in the standard Martini 3 database.

Here, we present refined Martini 3 parameters for 186 metabolites found in bacterial and eukaryotic cells^15,18^. They include updated nucleotide cofactors, flavins and phosphorylated sugars, cofactors which are crucial for biological functions. Following the Martini framework^1,5,19^, we used all-atom MD simulations as references to generate initial parameters that were subsequently refined based on numerical stability, and on properties such as solvent accessible surface area (SASA) and LogP measurements. The resulting Martini metabolome will allow high-throughput smaller-scale studies investigating individual metabolite interactions, alongside enabling large-scale simulations closer to *in vivo* conditions. We demonstrate the versatility of applications for this set of molecules with two showcases: the binding of ATP to an ABC transporter, and the permeation of glycerol across a lipid membrane.

## Results

### Selection of metabolites

To ensure a large coverage of biologically relevant metabolites, metabolites listed in a bacterial cell (JCVI-Syn3A)^15^ and from the human mitochondrial matrix^20^ were chosen for the Martini metabolome. The resulting 186 metabolites represent a broad range of chemical classes, including nucleotides, ions, carbohydrates, amino acid derivatives, lipids, amino acids and cofactors (Figure 1, Table S1). Combined with the already existing models, including molecular oxygen^21^ and those related to lipid metabolism^22^, the Martini 3 metabolome now covers 97.5% of the metabolites present in JCVI-Syn3A, and 98.7% of the mitochondrial matrix. Even though the selection was distinct from the database, the metabolite library covers the vast majority of the abundant metabolites measured in *E. coli*^18^.

**Figure 1.**
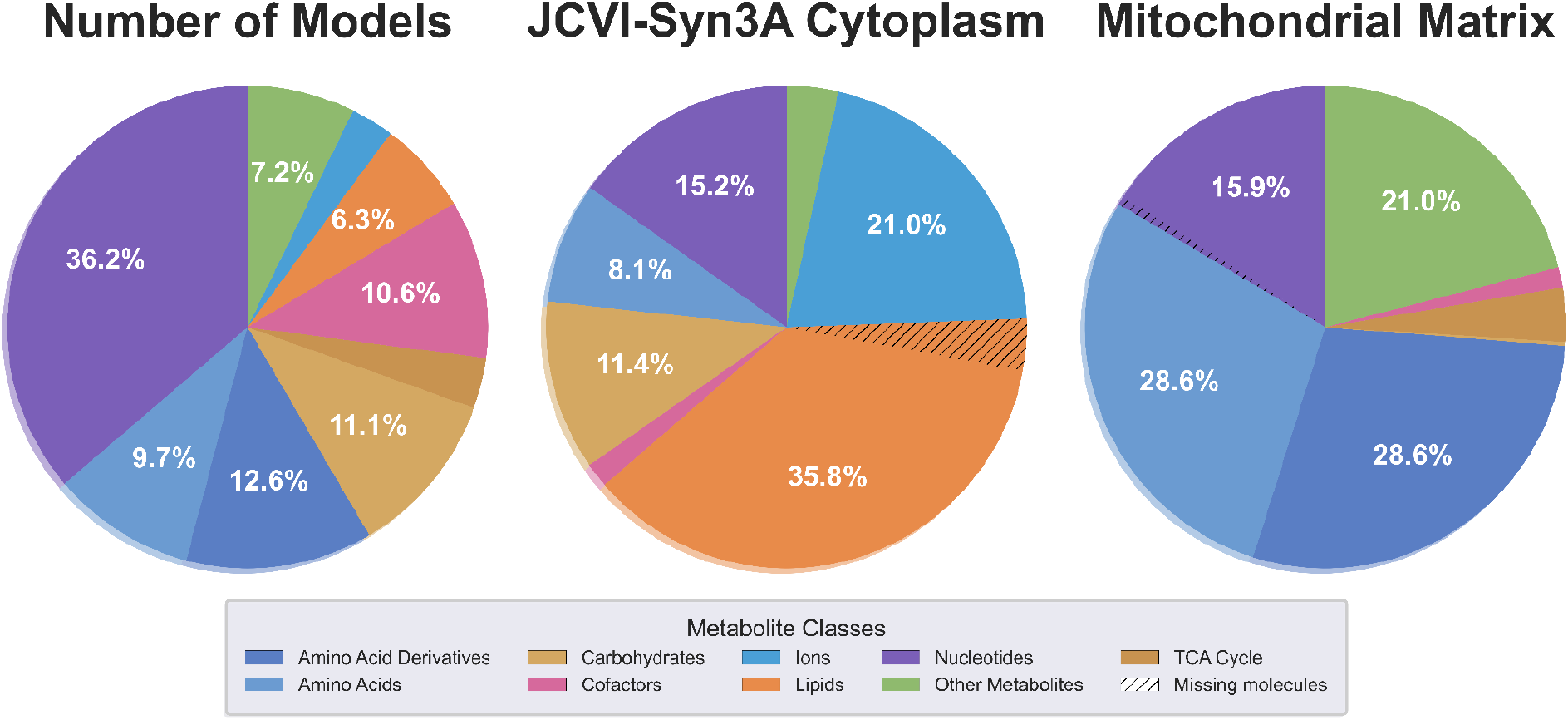
Overview of the parameterised metabolites. Left) Distribution by number of molecules parameterised in this study (186 in total), showing the diversity of newly available metabolites. Middle) Molar concentration distribution in the JCVI-Syn3A cell cytoplasm^15^. Right) Molar concentration distribution in the human mitochondrial matrix^20^. Metabolites are classified into functional categories (nucleotides, amino acids, amino acid derivatives, carbohydrates, TCA cycle intermediates, cofactors, lipids, ions, and other metabolites) and colored accordingly. In the middle and right panels, the hashed sections of segments represent molar concentrations in the respective metabolomes with models not yet available in Martini 3. Percentages are shown for classes representing ≥5% of the total.

The ionization state for each metabolite was assigned using SwissParam^23,24^ to identify the dominant species at pH 7. Alternative protonation states are also provided for some nucleotides (see Table S1) to capture species that might be desired by the simulation community.

### Parameterization strategy

The parameterisation strategy adhered to the general Martini guidelines. Following an atomistic simulation, a molecule was mapped to a pseudo-CG trajectory (Fig. 2A). CG molecular topologies were designed for each molecule to reproduce the respective bonded parameters of their pseudo-CG trajectory (Fig. 2B). Bead assignments were made by following standard mapping procedures, keeping chemically functional groups together. The bead type was initially assigned based on the partitioning behaviour of the underlying group. Thus, throughout the validation of non-bonded parameters, the Martini bead type for every molecule was assigned using the validated and suggested bead types for the force field^1,5,19^. Solvent accessible surface areas (SASAs) were assessed against all-atom references to ensure good volumetric matching^25^ (Fig. 2C-D). In addition to the usual parameterisation approach, we stress tested every molecule at high concentrations to further ensure numerical stability in crowded environments. The parameterisation protocol is described in full in the Method section. Topologies, and other parameterisation reference data for all the 186 metabolites parametrized in this study can be found in the GitHub repository https://github.com/Martini-Force-Field-Initiative/M3-Metabolome.

**Figure 2.**
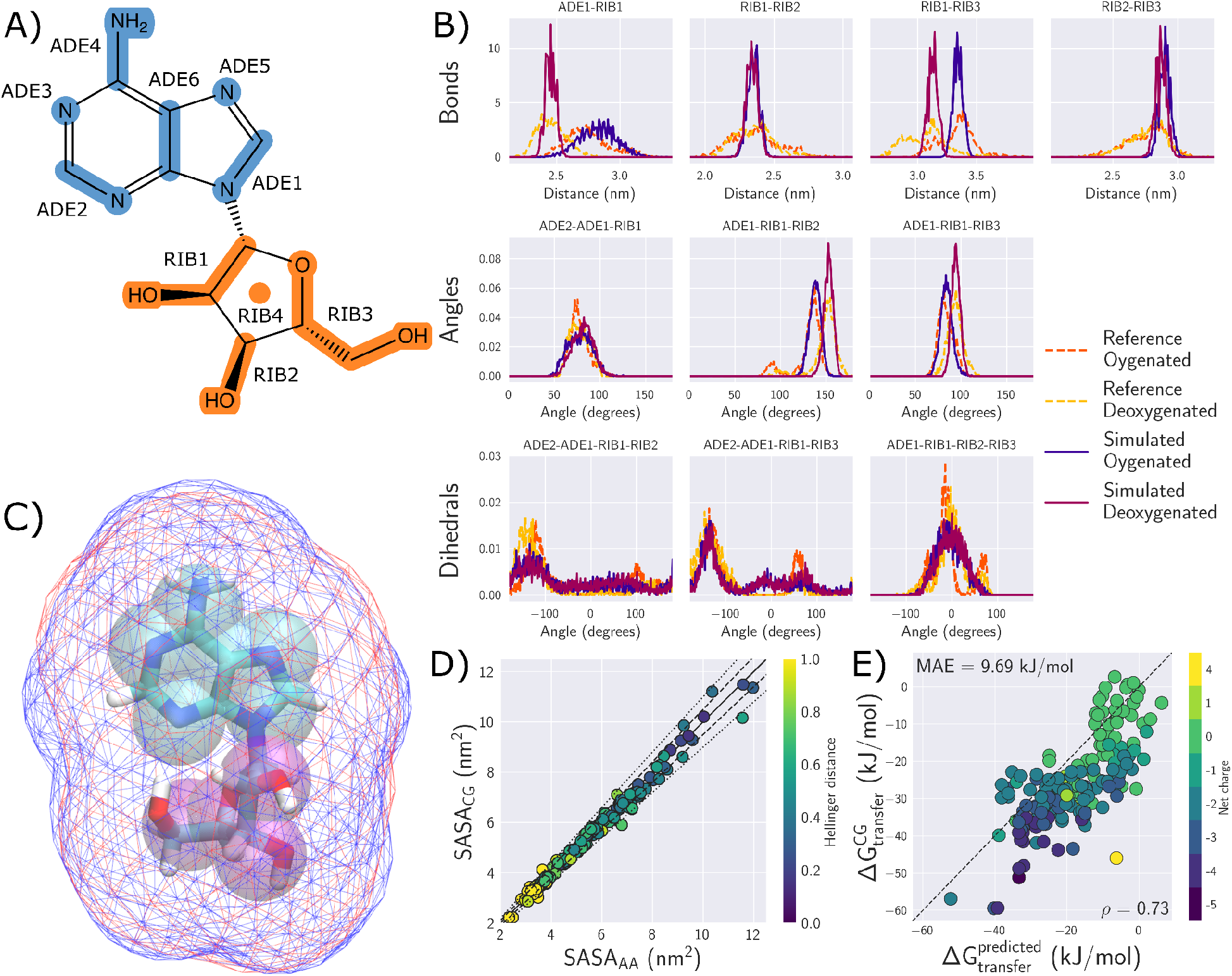
Parameterisation strategy of metabolites. A,B) Examples of molecular mapping and bonded interaction matching for adenosine. C) Connolly surfaces of adenosine at atomistic (blue) and CG (red) resolutions. D) SASA for all metabolites, coloured by the Hellinger distance between the atomistic and CG distributions. The dashed and dotted lines represent 5% and 10 % deviations respectively. E) Water-octanol partitioning free energy of transfer for metabolites, coloured by net charge. The mean absolute error and Spearman rank are indicated by MAE and ρ, respectively.

In general, the molecules described in this work fall into two categories: non-nucleotides, and nucleotides. The former class were generally parameterised *ab initio*, using the process described above. The latter class (and other associated molecules such as sugar-based metabolites) were parameterised using existing fragments in a self-consistent and hierarchical manner. For example, parameters for free nucleobases and sugars have already been developed for the Martini 3 force field^1,5^. Where available, common interactions were grouped together across molecules in order to ensure consistency across the set. For instance, the ADE2-ADE1-RIB1-RIB2 dihedral present in adenosine (Figure 2B) is also present in AMP, ADP, and ATP. Such interactions were therefore fitted once, against an average distribution, and optimised across all molecules where present. Bonded interaction validation figures are available for every molecule in the associated repository.

### Validation against partitioning data

One of the major targets of Martini parameterisation is the partitioning behaviour of molecules between water and other organic solvents. To this end, we computed the free energy of transfer between water and octanol (LogP) via alchemical transformations. The results can be found in Table S1. Ideally, it would be possible to compare the calculated partitioning free energies to an experimental reference, but this data is only available for a limited set of molecules. In Figure S1, we show that for molecules with experimental reference data, the calculated partitioning behaviour of the models matches closely, at least for neutral molecules.

To generate a comprehensive comparison dataset, we additionally used the SwissADME server^26^ to generate predicted values for all the molecules presented in this work using the XLOGP3 method. Figure 2E shows a comparison between the estimated partition coefficients of all the molecules and the predictions made by the server. Compared to the prediction, the transfer free energy of the metabolites has a mean absolute error (MAE) of 9.7 kJ/mol. While this does not represent a high degree of accuracy with respect to the predicted values, it is likely that the predicted values for many of these molecules (particularly those which are either polar or charged), does not represent an accurate prediction themselves.

In an effort to better understand the limitations of the predictors we compared XLOGP3 predictions to experimental LogP values used in recent Martini 3 parameterisation papers ^5,19,27–29^. As shown in Table S1 and Figure S2, XLOGP3 predictions are in good agreement with small molecule partitioning data. Additionally, LogP values for simple sugars as well as for a more complex molecule like riboflavin are close to the experimental values. However, and as we note in Figure 2E, predications start deviating strongly for charged molecules. Further, while for some sugars LogP values are predicted accurately, for other sugars they are not. Other LogP predictors generated by SwissADME do not lead to an overall improvement (Figure S3).

We therefore conclude that in general LogP predictors like XLOGP3 can be a good starting point if no LogP data is available. However with increasing molecule complexity predictions can be unreliable while for charged molecules a strong deviation is to be expected. As many molecules within the metabolome are charged, and in the absence of comprehensive experimental data, we therefore believe that the current estimates of partitioning behaviour for the Martini metabolome are faithful to the chemical nature of the underlying molecules.

### Showcases

#### ATP binding

The Martini 3 metabolome opens the way to study the interaction of metabolites with other biomolecules. To showcase these opportunities, we selected two examples of basic cellular importance, namely the protein binding of ATP and the membrane permeation of glycerol.

Due to its biological importance, the metabolite chosen to showcase protein-ligand interactions was ATP (Figure 3A). ATP was simulated with a protein from the ATP-binding cassette (ABC) transporter family, namely NosDFY, which is a nitrous oxide reductase^30^. The structure has been solved in an ATP-bound state, with the ligand bound to both copies of the nucleotide-binding domain (PDB ID: 7O17), which was used to build the CG starting state in our simulations. Note that an elastic network was used to retain the secondary structure and the tertiary fold of the protein. Three different systems were simulated: the whole transporter in a lipid bilayer bound to ATP, the soluble NosF dimer with ATP bound and the soluble dimer with ATP started in bulk solution (Figure S4). When the holo form of the whole complex was simulated, ATP was observed to stay within the binding site throughout five replicate simulations of 1 µs (Figure S5A).

**Figure 3.**
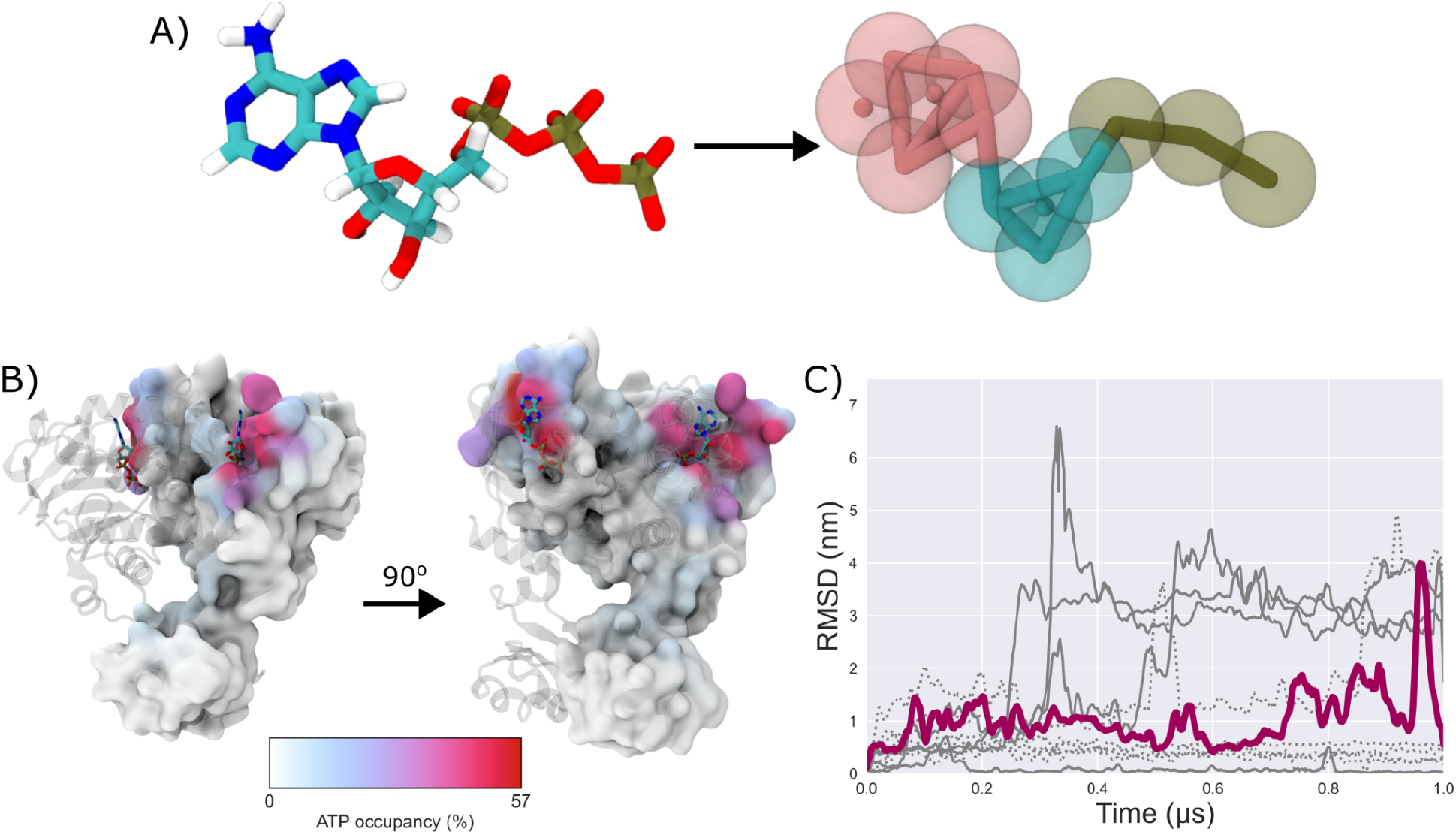
Metabolite interaction with proteins. A) A comparison of the atomistic representation of ATP (left) and the CG ATP (right). For the CG representation, the particles are shown as beads with bonds shown as sticks, and virtual sites as solid dots. B) The surface of the protein shows the ATP occupancy with NosF throughout simulations, with the darker color showing increased occupancy. The stick representation of ATP denotes the canonical binding sites from the cryo-EM structure. C) A plot showing the root mean squared deviation (RMSD) of each ATP (one with a solid line, the other with a dotted line) from the initial starting position.

To study possible entry and exit pathways, additional simulations were performed on just the soluble ATP-binding protein homodimer (NosF), again with ATP bound initially. Within the majority of simulation time, the ligand stays bound with occupancy of the binding site at ∼60% (Figure 3B). Interestingly, in one replica, an ATP leaves the known binding site, the molecule returns to the starting position after sampling the solvent (Figure 3C), showing that our ATP model captures the correct biological behaviour. To further test the ability of the molecule to find the binding site from the bulk, two copies of the ligand were placed in the solvent. Over a cumulative 5 µs simulation time, ATP was observed to spontaneously bind to the canonical binding site (Figure S5B).

#### Glycerol permeation of lipid membranes

The permeation of metabolites across lipid bilayers provides another test of parameter accuracy under biologically relevant conditions. Here, we investigated glycerol transport across the bacterial membrane of the JCVI-Syn3A minimal cell. Glycerol represents a particularly relevant case study for the minimal cell, where this essential metabolite must rely on passive diffusion due to the absence of a dedicated transporter in the reduced genome^31^.

We constructed a membrane model of the native Syn3A lipid composition containing cholesterol, sphingomyelin, cardiolipin, palmitoyl-oleoyl-phosphatidylcholine (POPC), and dioleoyl-phosphatidylglycerol (DOPG) at experimentally determined concentrations^15^ (Figure 4B). The potential of mean force (PMF) for glycerol translocation was calculated using umbrella sampling, and permeation rates were determined by combining the free energy profile with position-dependent diffusion coefficients calculated within each umbrella window (Figure 4C). Our simulations yielded a glycerol permeation rate of 7 ± 2 nm/s through the Syn3A membrane.

**Figure 4.**
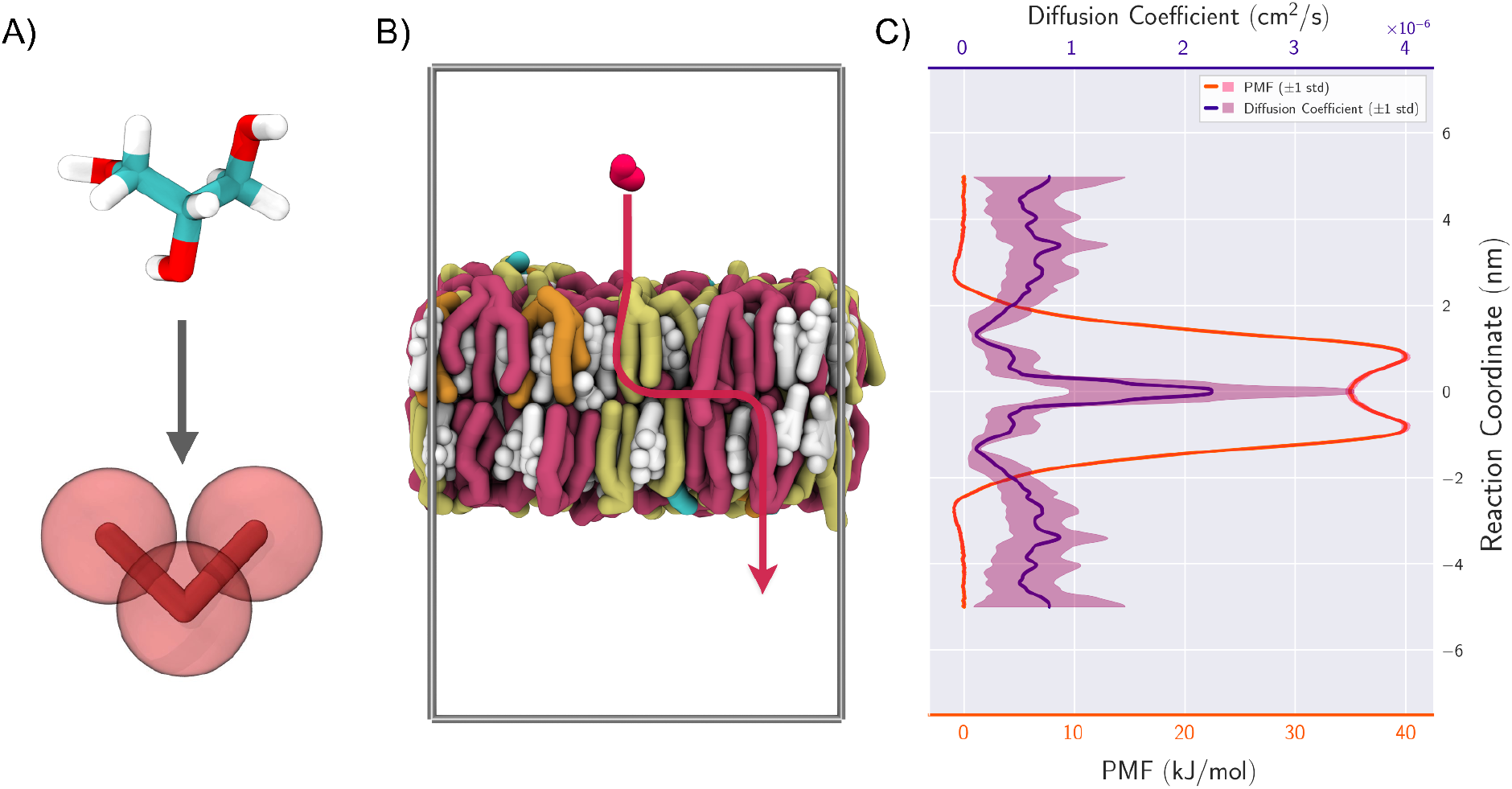
Glycerol permeation across a bacterial membrane. A) A comparison of the atomistic representation of glycerol (top) and the CG glycerol (bottom) B) Snapshot from umbrella sampling simulation showing glycerol (red) above the membrane containing cholesterol (white), cardiolipin (pink), POPC (orange), DOPG (cyan), and sphingomyelin (yellow). In red a representative permeation pathway through the membrane is shown. C) The calculated potential of mean force and diffusion profile for glycerol permeating through the JCVI-Syn3A membrane.

Given the generic speedup in dynamics inherent to CG models, we consider this value to be in good agreement with a recent experimental measurement of 3 ± 2 nm/s for glycerol permeation across a membrane of similar composition^32^. Noting that the permeation rate of a permeant is dominated by the barrier height of the PMF, the calculated permeation rate validates that our model parameters capture the free energy barrier needed for glycerol transport across the Syn3A membrane.

## Discussion

The metabolite library presented in this work addresses a fundamental limitation that has constrained the biological realism in molecular simulations targeting cellular environments with the Martini 3 forcefield. Without validated parameters spanning the chemical diversity of biological systems, researchers must either omit metabolites that are present in their experimental target systems or undertake laborious parameterization efforts that become prohibitive when attempting to capture realistic systems with high molecular diversity. While ongoing initiatives to centralize and distribute Martini parameters have improved accessibility, a significant lack of metabolite parameters has persisted^33^. By providing validated Martini 3 parameters for 186 cellular metabolites representing major chemical classes in bacterial and eukaryotic cells, this work opens the door to modeling in situ cellular environments that have previously been limited in complexity and realism. Current efforts are already underway to simulate the JCVI-Syn3A cytosol utilizing the Martini 3 metabolome.

While we have demonstrated the potential of the CG models in this work to already reproduce basic biophysical phenomena, we believe that the models still have significant potential for further improvement. Beyond numerical stability and basic physicochemical properties, we have not performed extensive validation testing for any molecule in the database. Future use of more targeted experimental reference data, along with additional considerations during parameterisation, could almost certainly result in more optimised models for specific applications.

Although our current library covers the major metabolite classes found in bacterial and eukaryotic cells, the framework developed here provides a foundation to fill in remaining gaps. For instance, the lipids class is slightly less well covered in comparison to the others, with metabolites such as galactose diacylglycerol (GAL-DAG) and CDP-diacylglycerol missing. The recent development of an extended Martini 3 lipidome offers further guidelines for their parametrization^22^. Similarly, parameters for CO_2_ have not been included or verified, although recent work on parameterising O_2_ for the Martini 3 force field offers a potential strategy for pursuing such work^21^.

Furthermore, this metabolite library has the potential to serve as a comprehensive dataset for the training and validation of machine learning approaches to automate CG topology generation^34,35^. Machine learning approaches for force field parameterization require diverse training data that spans the chemical space of biological molecules^36^. This metabolite library provides such a foundation, with validated Martini parameters across multiple chemical classes that represent the molecular diversity found in cellular environments.

The availability of these validated parameters transforms cellular metabolites from a technical barrier into a standard component of Martini modeling workflows. These parameters allow researchers to focus on fundamental biological questions, from metabolite-protein interactions to cellular crowding effects. As simulations increasingly approach biological complexity, this metabolite dataset provides essential components that bridge the gap between simplified model systems and the rich chemical environments found in cells.

## Methods

### Parameterisation strategy

Parameterisation followed the standard Martini 3 parameterisation pipeline^1^. First, atomistic simulations for each individual molecule were performed. The trajectories were then mapped according to a consensus mapping scheme to a coarse-grained resolution. The mapped trajectories were then analysed, such that CG molecular topologies could be generated. Once bonded distributions between resolutions were matched as optimally as possible, the molecules were further validated by comparisons of solvent accessible surface area (SASA), and the transfer free energy between octanol and water. Molecular topologies were further stress tested to ensure stability in crowded environments.

### Atomistic reference simulations

SMILES strings were compiled and uploaded to the swissparam server^23,24,37^ to obtain simulation input parameters and structures for the charmm36 force field. Simulations were conducted using Gromacs 2024.1. Single copies of molecules were placed in dodecahedral simulation boxes, 5 nm from the edge of the box, using the gromacs editconf program. The molecule was subsequently solvated with an ionic concentration of 0.15 M NaCl, plus additional ions for system neutralisation. The system was subsequently subject to short equilibration simulations in the NVT and NPT ensembles for 1 ns, before a 1 μs production simulation. Solvent-free trajectories are available as part of the supplementary information. For all simulations, the time step was 2 fs. Where pressure coupling was used, the pressure was set to 1.0 bar with a compressibility of 4.5×10^-5^ bar^-1^. Electrostatic interactions were treated with PME with a cut off of 1.2 nm. During equilibration the C-rescale barostat was used, and production simulations were performed using the Parrinello−Rahman barostat with isotropic pressure coupling^38,39^. Simulation temperature was controlled with the V-rescale thermostat^40^.

### Coarse-grained simulations

Unless stated otherwise, all CG simulations in this work followed the standard Martini 3 protocol described below. Simulations were conducted using Gromacs 2024.1. Simulations of single molecules were prepared directly from the solvent-free mapped atomistic systems, using the same simulation cell dimension. Molecules were then solvated, and charge neutralised with 0.15 M NaCl. The system subsequently underwent energy minimization using Gromacs’ steepest descent algorithm, followed by equilibration and production simulations. Electrostatics were treated with reaction-field with a cutoff of 1.1 nm, and van der Waals interactions were treated with potential-shift-verlet, also with a cutoff of 1.1 nm^41^. Additionally, we used the recent recommended settings from Kim et al. (verlet − buffer − tolerance = −1, rlist = 1.35 nm) to correct neighbourlist artifacts^42^. Pressure was maintained isotropically at 1 bar using the c-rescale barostat during equilibration, and for production, the Parrinello−Rahman barostat. Compressibility was set to 3.0×10^-4^ bar^-1^. The V-rescale thermostat was used to maintain temperature at 300 K for all simulations. A time step of 10 fs was used for equilibration, which was increased to 20 fs for production runs.

### Mapping

Molecular mapping was conducted according to the standard Martini 3 mapping rules. The mapping of bonded distributions was performed using the fast forward Python package, and where necessary collated by custom scripts^43^. Mapping (.map) files following the Vermouth format were generated using the cgbuilder tool^44^, and are supplied in the supplementary information. For future cheminformatical purposes, we also provide the CGsmiles strings for mapping information^45^.

### Bonded interactions

Initial topologies of molecules were generated using the ff_inter program of the fast forward Python package^43^. As fast_forward is designed to generate high-fidelity bonded parameters based on mapped simulations, in the first instance these parameters were tested without further inspection. Initial conformers of molecules were generated using Polyply^46^, and solvated in a 5×5×5 nm simulation cell with 0.15 M NaCl, and neutralising counter ions. The systems were then subjected to steepest descent energy minimisation, followed by equilibration and production simulations of lengths 10 ns and 500 ns respectively.

Following an initial simulation of a molecule, the resultant bonded distributions were validated against their mapped references using the ff_assess program of fast_forward. ff_assess scores simulated bonded distributions against the references used during fitting, and returns an average Hellinger distance between the two probability distributions. A score below 0.3 is considered a good agreement with an atomistic reference for a molecule, which was used as a target throughout parameterisation. Molecular topologies were then optimised according to which bonded parameters were in least agreement with their respective references. This was achieved by several methods, such as varying force constants, or adding additional exclusions between beads. In the case of dihedral terms, initial fits using ff_inter may have overfit the mapped distribution and caused instabilities. In this case, the ff_inter program was rerun with a smaller value for the maximum multiplicity of dihedral to be used during fitting, to achieve better stability at the sacrifice of a degree of fidelity.

### Stress testing

To ensure CG model stability, once the bonded parameters of a single copy of a molecule had been validated, 50 copies were simulated in a 10×10×10 nm cubic box for 500 ns. If the simulation proceeded without error, then the model was determined to be stable for release. If the simulation did not run to completion, the model was reviewed for stability by repeating the mapping and interaction optimization steps described above as necessary. The simulation input parameters followed the standard Martini ones described above.

### Non-bonded validation

Partition coefficients (LogP) for each molecule was calculated using alchemical free energy transformations. The free energy of solvation was estimated in both water and hydrated (0.3 M water) octanol^19^, using 21 steps during which the non-bonded interactions between the solute and solvent were turned off. Soft core potentials were used using the recommended (sc-alpha = 0.5, sc-power = 1) values, along with a stochastic dynamics integrator^47^ and an inverse friction constant of 1.0 ps. The transfer free energy (ΔG) was calculated using MBAR^48^ as implemented in the alchemlyb python package^49^.

### Showcase 1: Ligand binding

The atomistic structure file for NosDFY^30^ was downloaded from the PDB database. The protein was converted to a Martini representation using martinize2^50^ with an elastic network (merging all chains when relevant) using a force constant of 700 kJ/(mol nm^2^). The elastic network was applied in the range of 0.5-1 nm. When the metabolite was started within the binding site, the coordinates were obtained by mapping with CGBuilder^44^. When the ligand was started outside of the binding site, the *gmx insert-molecules* tool was used to generate independent starting positions for two molecules. The whole NosDFY transporter was embedded in a membrane (POPE:POPG, 4:1) using *insane*^51^. Systems were solvated and 0.15 M NaCl added, with neutralising counter ions. The systems were energy-minimised, equilibrated and production simulations performed using the standard Martini protocol described above. For the simulations without a bilayer, isotropic pressure coupling was used at 1 bar. For the entire protein complex in a bilayer, semi-isotropic pressure coupling was used at the same reference pressure. All simulations were performed at 310 K. Each system was simulated for 5 x 1 µs.

Occupancy analysis was performed using PyLipID^52^ and visualised using VMD^53^. RMSD of the ligand was calculated using *gmx rmsd* relative to the protein backbone and plotted using Matplotlib^54^. Visualisation of protein structures was aided by the MartiniGlass package^55^.

### Showcase 2: Membrane permeability

A membrane model was constructed to reflect the native JCVI-Syn3A envelope composition: cholesterol (59%), sphingomyelin (18%), cardiolipin (17%), linked POPC (4%), and phosphatidylglycerol (2%)^15^. The bilayer was assembled using the insane tool in a 10 x 10 x 20 nm^3^ simulation box^51^. The system was then solvated with Martini water, where a subset of water beads were replaced with ions to neutralise the model and add an ionic strength of 0.15 M NaCl.

The system was equilibrated and simulated using the standard Martini3 simulation protocol as described previously, with the addition of a semi-isotropic pressure coupling. The bilayer was equilibrated for 200 ns at 300K. After equilibration, umbrella sampling was performed with glycerol restrained at intervals of 0.1 nm from the bilayer center until 5 nm into the aqueous phase. Harmonic restraints with a force constant of 1000 kJ mol^-1^ were applied to the glycerol center of mass. Each umbrella window was simulated for 100 ns. The potential of mean force was reconstructed using WHAM with uncertainties estimated from 500 bootstrap iterations (Figure S6A).

Position-dependent diffusion coefficients along the transmembrane axis were calculated using the method described by Hummer^56,57^. For each umbrella window, five replicas of 10ns were performed with the glycerol center of mass saved at every timestep. These trajectories were used to calculate the local diffusion coefficient according to:

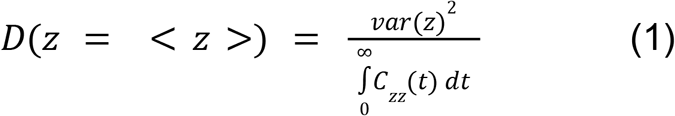

where *C*_zz_ (*t*) is the autocorrelation function of the reaction coordinate (Figure S6B). Due to the high uncertainty in the calculated diffusion coefficients, a moving average filter with a window size of 2 was applied to the mean diffusion profile.

The membrane permeability was then calculated by integrating the position-dependent resistance across the membrane using the relation^58^:

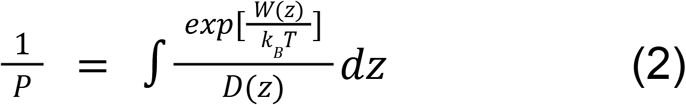

where W(z) is the potential of mean force, k_B_ is the Boltzmann constant, T is the temperature. To propagate uncertainties into the permeability estimate, this calculation was performed 100,000 times using randomly sampled combinations of the replicate diffusion profiles and bootstrapped PMF profiles (Figure S6C). This bootstrap analysis resulted in a permeability coefficient of 6.81 ± 1.74 nm/s.

## Data Availability

Full supporting information is available on Zenodo at 10.5281/zenodo.18622430. A subset of supporting information (CG topology and coordinate files, mapping files and bonded distribution comparisons) are available on the M3-Metabolome repository on GitHub: https://github.com/Martini-Force-Field-Initiative/M3-Metabolome

## Supporting information

Figures for additional LogP validation (S1-S3), details of ATP binding simulation setups (S4, S5), and additional validation for glycerol permeation (S6) are available in the supplementary information (PDF). Table S1, containing a full list of molecules parameterised along with additional details (LogP values, SMILES strings used, CGSmiles descriptors, notes on protonation states), is supplied as a separate csv file.

## Acknowledgements

S.J.M. acknowledges funding from NWO through the Summit Grant “Evolving life from non-life (EVOLF)”. J.A.S. and S.J.M acknowledge funding from NWO through the NWA grant “The limits to growth: The challenge to dissipate energy”. C.M.B., L.G., and S.J.M acknowledge funding from the ERC with the Advanced grant “COMP-O-CELL”. C.B. and S.J.M. are supported by the Novo Nordisk Foundation Grant NNF20OC0063808, “BOUNDLESS”. We would like to thank SURF for access to Snellius computing resources. We thank Fabian Grünewald for assistance with preparing CGsmiles strings for molecular mappings.

