## Supplementary material for "The Martini 3 Metabolome": SI

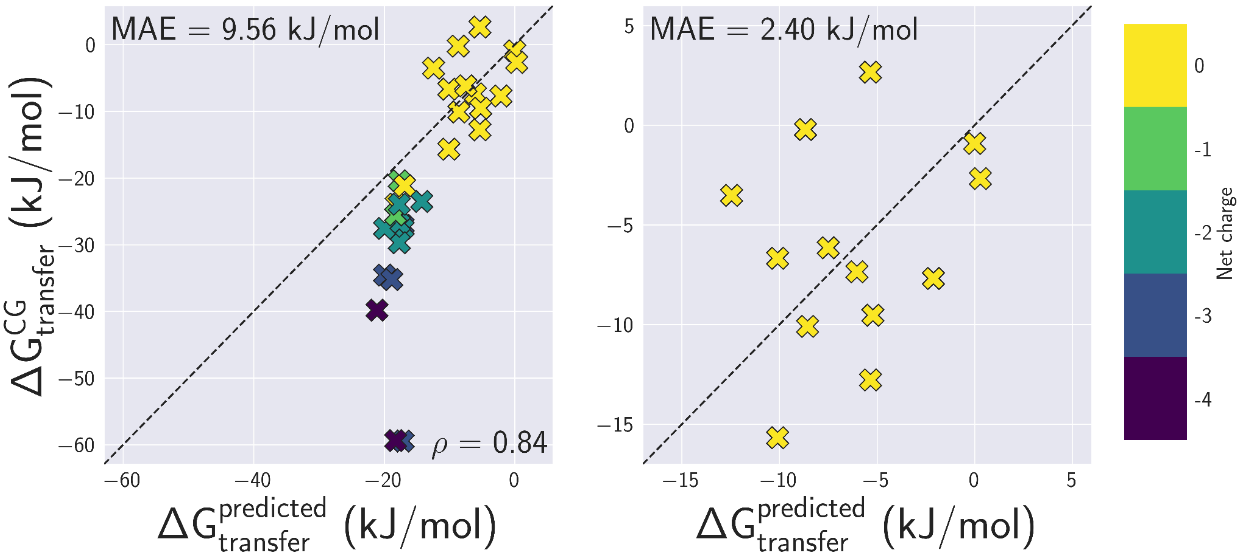


**Figure S1. LogP comparisons with experimentally available data.** Left, comparisons with all experimentally available data. Right, comparisons with only neutral molecules.

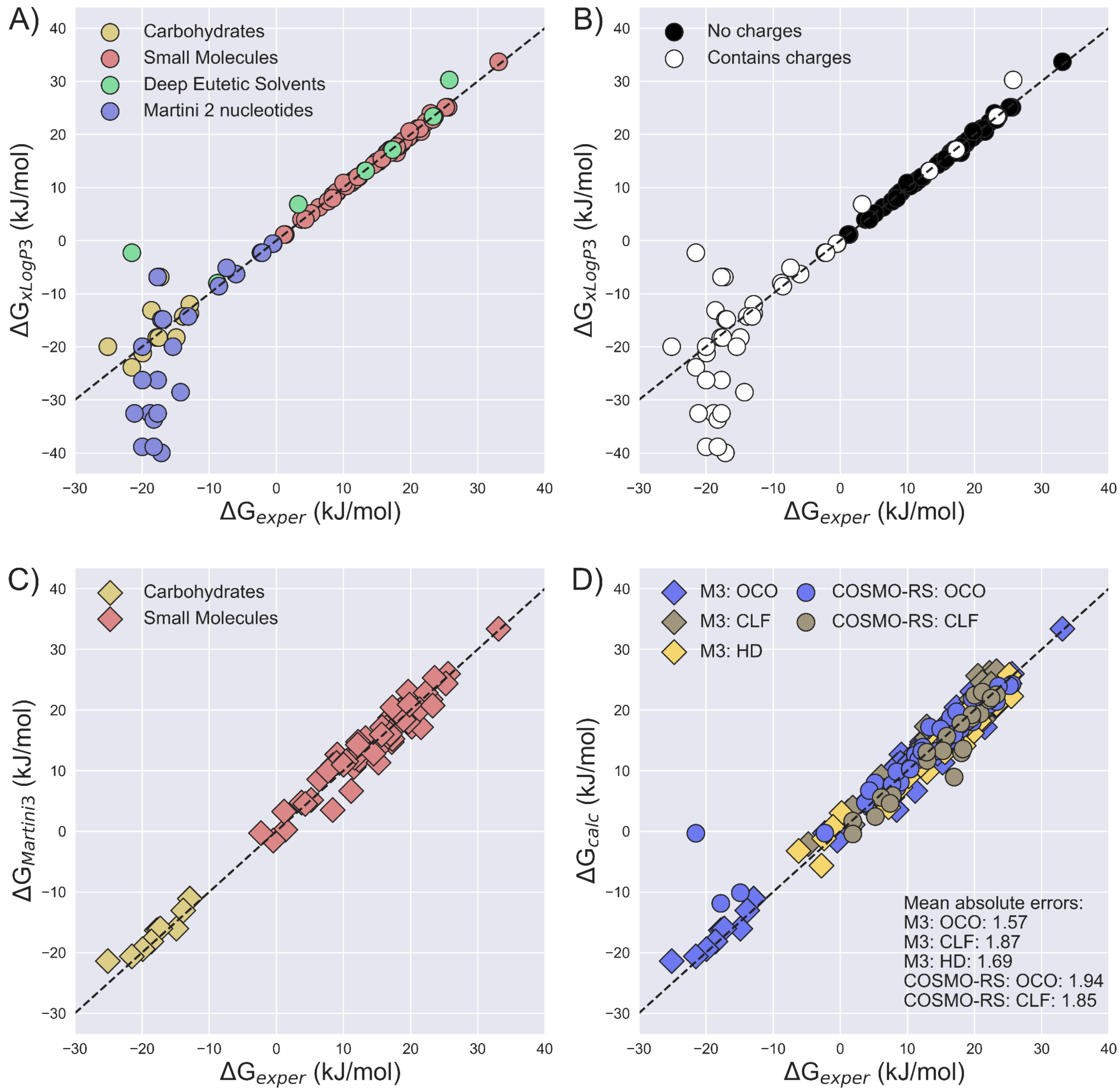


***Figure S2. Using LogP predictors to estimate partitioning.*** *A,B) experimental ΔG (water-octanol) values plotted for four classes of molecules with known data for Martini force fields, and their xLogP3 prediction. In A) the class of each molecule is indicated by colour, and in B), the presence of charges. Note that the 'contains charges' class includes zwitterionic molecules. C) The experimental value of ΔG is plotted against the Martini 3 partitioning data for carbohydrates and small molecules. D) Comparison of experimental data to partitioning predictions using COSMO-RS for water-octanol and water-chloroform for the small molecules and carbohydrates, and calculated using Martini 3 models. Hexadecane partitioning is not available in COSMO-RS, but the Martini 3 data for small molecules is included for reference.*

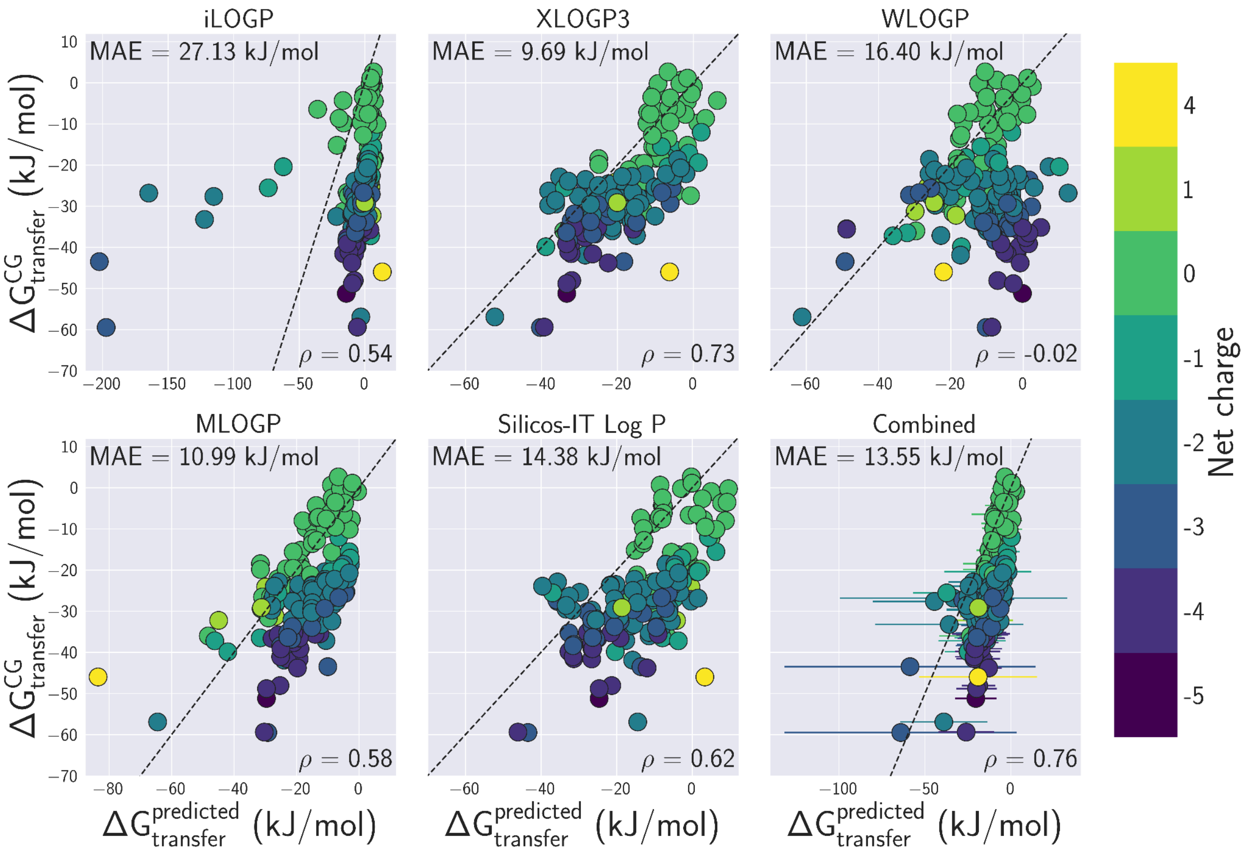


***Figure S3.*** *Comparison of computed ΔG values (y axis) to the five different predictors used by swissADME. The bottom right panel ('combined') shows the values averaged across all the predictors. Dashed lines indicate perfect agreement between prediction and measured ΔG.*

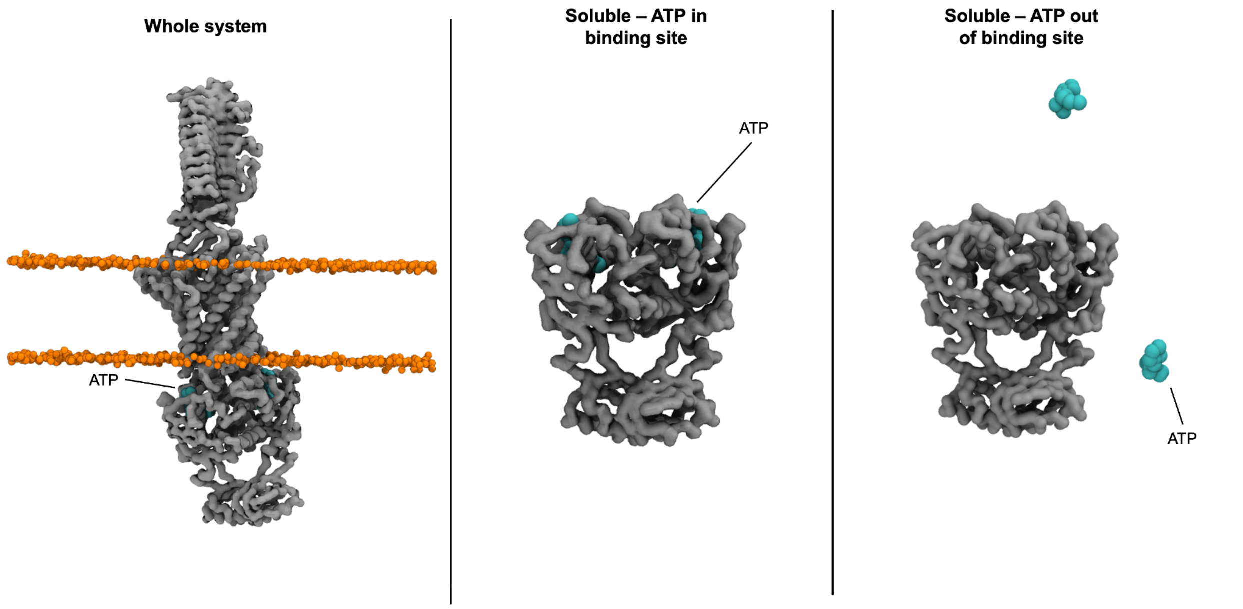


***Figure S4:*** *Starting poses of the NosDFY systems simulated with ATP. Whole transporter embedded in a lipid bilayer with ATP bound (left). NosF, the soluble nucleotide-binding domain, dimer with ATP bound (middle). NosF, the soluble nucleotide-binding domain, dimer with ATP in bulk solution (right). The protein is shown as a gray surface, the ATP is shown as cyan spheres and the lipid phosphate are shown as orange spheres.*

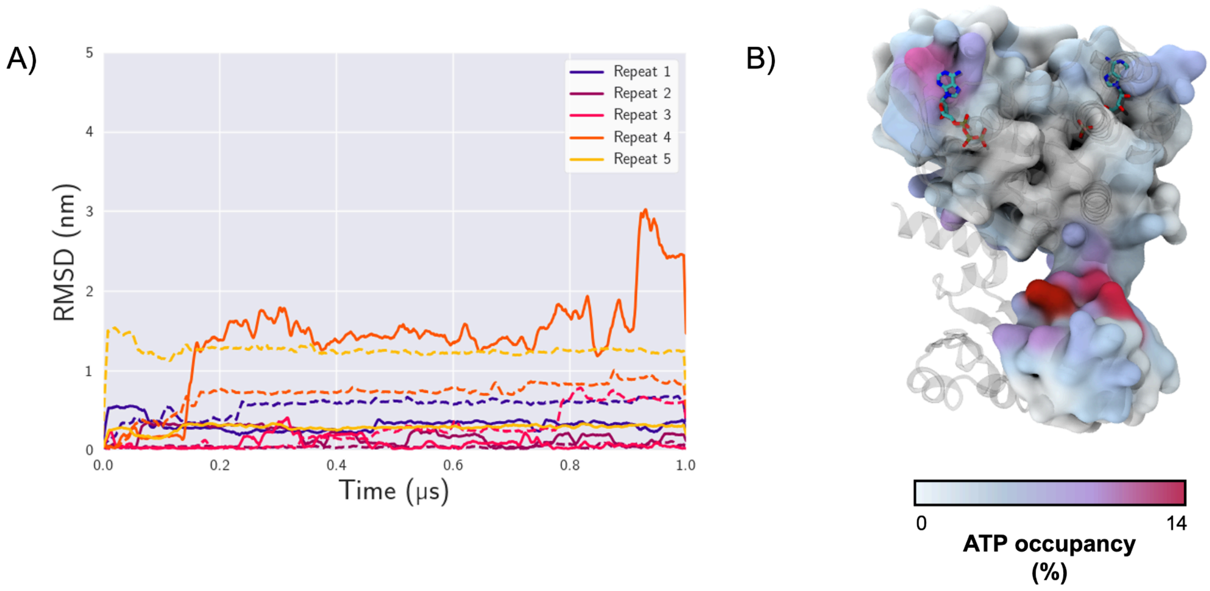


***Figure S5:*** *ATP interaction with NosDFY. A) A plot showing the root mean squared deviation (RMSD) of each ATP (one with a solid line, the other with a dashed line) from the initial starting position with the entire complex embedded in the membrane. The data is averaged over 10 data points. B) The surface of the protein shows the ATP occupancy with NosF throughout simulations when the ligand was started in the solvent, with the darker color showing increased occupancy. The stick representation of ATP denotes the canonical binding sites from the cryo-EM structure.*

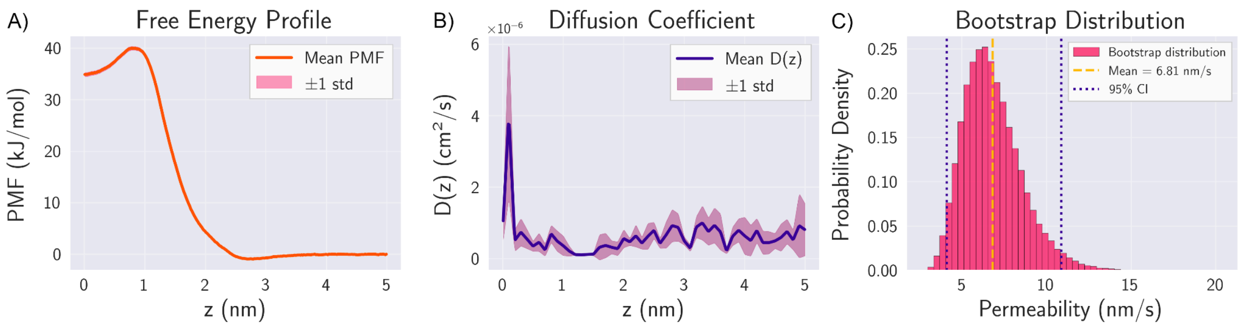


***Figure S6***: *Bootstrap analysis of glycerol permeability across the membrane. A) Mean PMF profile from 100 GROMACS bootstrap samples with standard deviation shown as the shaded region. B) Mean position-dependent diffusion coefficient from 5 independent replicates with standard deviation shown as the shaded region. C) Bootstrap distribution of permeability coefficients (n = 100,000 samples) calculated using Equation X. The mean permeability (yellow dashed line) and 95% confidence interval (blue dotted lines) are indicated.*
